# A high-throughput and sensitive method for food preference assays in the Argentine ant

**DOI:** 10.1101/2024.04.10.588882

**Authors:** Thomas Wagner, Henrique Galante, Tomer J. Czaczkes

## Abstract

Insects pose significant challenges in both pest management and ecological conservation. Often, the most effective strategy is employing toxicant-laced baits, which must also be designed to specifically attract and be preferred by the targeted species for optimal species-specific effectiveness. However, traditional methods for measuring bait preference are either non-comparative, meaning that most animals only ever taste one bait, or suffer from methodological or conceptual limitations. Here we demonstrate the value of direct comparison food preference assays using the invasive and pest ant *Linepithema humile* (Mayr, 1868) as a model. We compare the food preference sensitivity of non-comparative (one visit to a food source) and sequential comparative (visiting one type of food then another) assays at detecting low levels of aversive quinine in sucrose solution. We then introduce and test a novel dual-choice feeder method for simultaneous comparative evaluation of bait preferences, testing its effectiveness in discerning between foods with varying quinine or sucrose levels. While the non-sequential assay could not detect aversion to 1.25mM quinine in 1M sucrose, the sequential comparative approach detected aversion to quinine levels as low as 0.94mM. The novel dual feeder method approach could detect aversion to quinine levels as low as 0.31mM, and also preference for 1M sucrose over 0.75M sucrose. The dual-feeder method combines the sensitivity of comparative evaluation with high throughput, ease of use, and avoidance of interpretational issues. This innovative approach offers a promising tool for rapid and effective testing of bait solutions, contributing to the development of targeted control strategies. The method could also be easily extended to other ant species.

## Introduction

Ants are a major control concern: Pest ants pose significant economic threats due to their ability to infest agricultural crops, damage property, contaminate healthcare facilities, and contaminate stored food.^1,2^ Invasive ants threaten biodiversity, disrupting ecosystem processes, and can become significant ecological problems themselves.^3,4^

One of the best current approaches for eradication or control of ants is to use baits with a slow-acting toxicant. This method often surpasses traditional spraying because it targets the entire colony, including the queen and brood, by allowing ant workers to carry the toxicant into the nest.^5,6^ Baits have been shown to be effective both in the field and in buildings. ^7–10^

However, controlling ants poses considerable challenges: two-thirds of eradication efforts targeting invasive ants have proven unsuccessful.^6^ Alongside active bait abandonment or aversion^11,12^ a significant challenge lies in the availability of high-quality natural food sources, which ants often prefer, resulting in inconsistent bait consumption.^13,14^ To effectively address this challenge, it is crucial to understand the feeding preferences of ants. This information is vital for designing baits that effectively attract, and are consumed by, ants - particularly when they contain various ingredients such as toxicants, bittering agents, co-formulants, and attractors. If the bait, its additives, or its co-formulants do not align with the ants’ preferences, it may be rejected, or dispreferred compared to other available food sources. Testing the acceptance of baits and the various co-formulants they contain is thus crucial.

Commonly used methods for determining food preference include presenting foods and baits simultaneously, known as cafeteria tests, or sequentially offering solutions, often associated with odour stimuli. Preference testing is then conducted by measuring the number of ants attracted to various food sources or their associated odours, or by quantifying the amount of food consumed.^15^

Cafeteria-style experiments serve as valuable tools for understanding preferences in various species. For example, in *Apis mellifera* (Linnaeus, 1758), such experiments have revealed a preference for a sucrose-ethanol solution over a pure sucrose solution in dual-choice feeding situations.^16^ Similarly, the method was used to demonstrate that ants favour basic sugars combined with attractants, ^17^ and a preference for nutrients they are deficient in.^18^ Cafeteria-style experiments have also been used to examine how seasonal variation influences preference.^19,20^ In more applied settings, this approach was used to explore preferences for liquid versus gel baits, as well as for wet versus dry bait attractants,^21,22^ and investigate bait particle size preference.^23^

The other commonly used method for testing ant preference is the sequential-choice test, in which individual ants are presented with a series of choices sequentially to determine their preferences, either directly or by associating each option with an odour. For instance, sequential-choice tests have demonstrated that ants exhibit a preference for the first odour encountered,^24^ food sources without other nestmates around them,^25^ or closer food sources. ^26^ Preference can be influenced by various factors such as personal expectation, experience, social learning, and the timing of stimulus exposure. ^19,24,27–29^ Importantly, evaluation by comparison, where an individual’s evaluation of a food source depends on how it compares to other known options, interacts with the baseline evaluation of the food to shape feeding decisions.

To investigate comparative valuation, animals are usually trained to expect a certain quality or quantity of reward, which is then abruptly increased (positive incentive contrast) or decreased (negative incentive contrast).^29–34^ For instance, in a sequential preference test assay, *Lasius niger* (Linnaeus, 1758) ants were to evaluate and accept a fixed-quality sucrose solution differently depending on their previous experiences:^29^ During training, ants encountered sucrose solution droplets of either low or high concentration. Then, during testing, all ants encountered a sucrose solution of medium concentration. Ants expecting low-quality food exhibited greater acceptance of medium-quality food, while those expecting high-quality food showed the opposite pattern. Further experiments suggested that these contrast effects stem from cognitive rather than mere sensory factors. Food received within the nest from foraging ants influenced the perceived value of food encountered outside the nest.^29^ These findings underscore the importance of comparative evaluation in decision-making processes. When it comes to preference testing and the development of novel baiting strategies, it is essential to address the limitations of traditional methods. For instance, the common practice of marking individuals,^29,31,35^ required in order to test the trained individual ant in a sequential setup, can unintentionally induce stress and subsequently alter their preference. Preference can also be altered depending on which choice was presented first, or if the presented choice had an unexpected flavour.^24,36^ Presenting different choices simultaneously, but in different locations (as in cafeteria tests), can be vulnerable to side and location biases. Most critically, however, the cafeteria test approach prevents animals from making direct comparisons in real-time, thus potentially reducing their sensitivity. In response to these challenges, we have developed a new non-disruptive solution: the dual-choice feeder.

Here, we first demonstrate that comparative evaluation tests are much more sensitive than non-comparative tests for detecting a distasteful additive (quinine) in food. We then conduct a comparison between conventional sequential preference tests and our new dual-feeder method, using *Linepithema humile* as a model species. We test the ability of the dual-feeder method to differentiate between foods with added distasteful substances (quinine) and between foods with lower levels of attractive substances (sucrose).

## Materials & Methods

### Colony Fragments Maintenance

*Linepithema humile* (Mayr, 1868) were collected from Girona, Spain in April 2021. Ants were split into colonies, each consisting of one or more queens and 300 to 1000 workers. The colonies were housed in plastic foraging boxes (32.5 cm x 22.2 cm x 11.4 cm), with plaster of Paris on the bottom and PTFE-coated walls to prevent escape. Each box contained several 15 ml plastic tubes covered with red transparent plastic slide, partially filled with water, and plugged with cotton, acting as nests.

Colonies were maintained under a 12:12 light:dark cycle at room temperature (21- 25°C). Ants had continuous access to water via both the plugged tubes and a water feeder. Colonies were offered *ad libitum* 0.5M sucrose solution and freeze-killed *Drosophila melanogaster* (Meigen, 1830), but were deprived of food for 4 days prior to testing.

### Experiment 1 – Quantifying food acceptance using the established sequential testing method

The aim of this experiment was to test if ants with an expectation for pure sucrose solution show a reduced acceptance of solutions with small amounts of quinine. This is important, as before the dual-feeder can be tested and evaluated, an established method for testing comparative food acceptance must be demonstrated. This approach relies on comparing two groups of ants: naïve ants, and ants with recent previous experience of a comparator food source.

To quantify food acceptance by naive ants, a single ant was allowed access to a paper-covered 1x10cm linear runway via a drawbridge, following Wendt et al.^29^ The ant encounters a droplet of test solution at the end of the runway, and its behaviour was quantified. Initial acceptance was defined as sustained contact between the ant’s mandibles and the sugar droplet for ten seconds immediately after encountering the sucrose droplet. A short 3- or 10-second acceptance criterion is a commonly used method for assessing initial acceptance:^29^ 1M sucrose is consistently accepted within 10 seconds, while a lower concentration of 0.5M sucrose often leads to initial feeding interruptions. However, after an initial interruption, ants generally continued drinking the 0.5M solution until they are satiated, showing final acceptance. To quantify this acceptance criterion, in pilot studies we found that the average total drinking time of 1M sucrose in *L. humile* is approximately 90 seconds (Wagner et al., unpublished data). Thus, after 90 seconds, we noted whether the antś abdomen was visibly distended, an indicator of food ingestions, hence final acceptance.

Once an ant had been tested, it was removed from the platform and separated from the colony. Subsequently, both the paper cover and the sample solution were replaced. Various solutions with different sucrose-to-quinine concentration ratios were tested, including a control solution with 1M sucrose, and solutions with decreasing quinine concentrations (1.25mM quinine + 1M sucrose, 0.94mM quinine + 1M sucrose, 0.625mM quinine + 1M sucrose, and 0.3125mM quinine + 1M sucrose). These quinine levels were selected based on pilot studies in which *L. humile* showed high acceptance of a 1.25 mM quinine + 1M sucrose solution but consistently rejected higher concentrations in a no-choice assay. Thus, 1.25 mM quinine was designated as the maximum acceptable bitterness level. We then chose the lower concentrations based on serial dilution, as a good method for locating the aversion inflection point. In total, 143 ants from seven different colonies were tested over the five test treatments.

To qualify acceptance of experienced ants - thus comparative evaluation - individual ants stemming from a ‘donor’ colony (home colony) were given access to a pure 1M sucrose droplet at the end of the straight runway. While drinking, the ant was marked with a dot of paint on the abdomen and was allowed to return to a ‘recipient’ colony nest (related colony). This was done to ensure that food was not introduced to the ‘donor’ colony, which could potentially change the expectation of the ants which would be tested next. While the ant was in the recipient nest, the pure sucrose droplet was replaced with one of the sucrose/quinine solutions. The ant subsequently re-entered the straight runway via the drawbridge and was allowed to taste and feed on the test solution. In total, 199 ants from five different colonies were tested.

### Experiment 2 – Quantifying ant acceptance using the dual-feeder method

The goal of this experiment was to validate the findings of the initial experiment while demonstrating the benefits of this novel dual-feeder method.

We developed a new feeder design, which was 3D-printed using a stereolithography resin printer (Prusa SL1, Prague). 3D model files are available at https://doi.org/10.5281/zenodo.10953784. The feeder consists of two triangular wells which narrow into two parallel 0.35mm-wide channels, directing liquids to their tips through capillary action. The channels are separated by a narrow (0.30mm) gap (see Figure 1). This ensures that, regardless of which solution is contacted first, the ant will almost immediately come into contact with the alternative solution by touching it with its antennae, enabling an informed choice between the two solutions. Access to the feeder area is via a narrow bridge, ensuring that the ant is funnelled to the centre of the dual-feeder. Access to the dual-feeder is via a larger ‘landing platform’ – a raised platform surrounded by a water moat, and covered by a 1 x 10cm linear paper overlay, tapering to 2mm at the tip. Ants are allowed to walk onto a piece of paper in the nest, and then allowed to walk off the paper onto the landing platform paper overlay. From there, the ant walks without interference until it reaches the dual-feeder.

**Figure 1:**
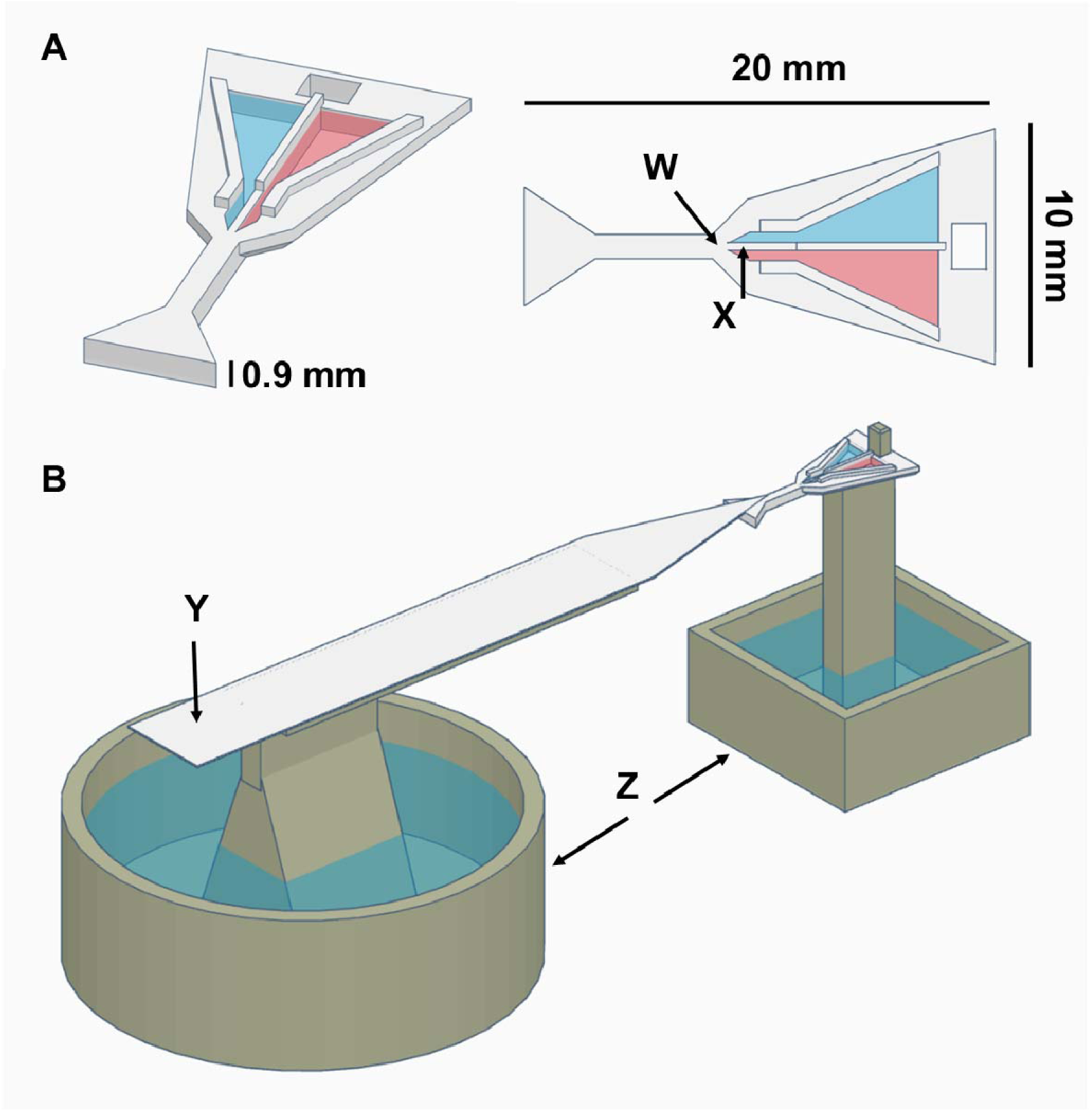
A: 3D model of the Dual-Feeder, W: Optimal drinking position, X: 0.30mm gap separation between the two wells containing solution 1 and 2 (blue and red coloured), B: Whole experimental setup display showing the dual-feeder connected to the 10 x 1 cm landing platform Y and attached to a pillar Z, surrounded by a water moat to prevent ants from escaping. 3D-model files for all printed parts are provided at https://doi.org/10.5281/zenodo.10953784.

#### Avoidance of quinine-containing solutions

In this experiment, one of the wells in the dual-feeder contained a comparator (here 1M sucrose solution) and a the other a test sucrose-quinine solution (or a control). These are presented simultaneously. Quinine in the test solution varied in concentration: 0.63mM, 0.31mM, and 0.16mM, all in a 1M sucrose solution. The side containing the test solution was systematically varied between ants. Ants encountered one of the drinking channels first either with their mandibles or antennae and began drinking. The initially encountered side and drinking time for each solution were recorded. In total, 179 ants from eight different colonies were tested.

#### Preference for higher molarity sucrose solution

Here, test solutions were of equal or lower sucrose concentration to the comparator, and contained no quinine. The following comparisons were tested: 1M vs 0.5M, 0.75M vs 0.5M, and 1M vs 1M. In total, 422 ants from nine different colony fragments were tested.

#### Statistical analysis

The complete statistical analysis output, and the entire dataset on which this analysis is based, is available from https://doi.org/10.5281/zenodo.10953784. Additionally, the post-hoc lateralisation analysis can be accessed at https://zenodo.org/records/14204095.

All graphics and statistical analysis were generated using R version 4.2.2^37^ and ggplot2.^38^ Sequential test data (Experiment 1) was analysed using binomial generalized linear mixed-effect models (GLMM).^39^ DHARMa^40^ was used to assess linear model assumptions and MuMIn^41^ to obtain a measure of goodness of fit. Dual-feeder proportion data (Experiment 2) was analysed using beta regressions.^42^ Analysis of variance tables were used to test the effects of regression coefficients.^43^ Estimated marginal means and contrasts were obtained using the emmeans package^44^ with Bonferroni adjusted values accounting for multiple testing.

## Results

### Experiment 1 – Quantifying food acceptance using the established sequential testing method

#### Bitterness acceptance comparison between naive ants and experienced ants

We compared the acceptance rates of naive ants, which were exclusively exposed to the sucrose-quinine mix, with experienced ants, which first encountered a pure sucrose solution before being presented with the sucrose-quinine mix. In terms of initial (first 10 second) acceptance scores, experienced ants showed significantly lower food acceptance than naive ants, showing a 70% lower acceptance for 1.25mM quinine (GLMM: z = 4.93, p < 0.0001) and a 54% lower acceptance for 0.94mM quinine (GLMM: z = 3.66, p = 0.0012) (see Figure 2). However, no difference in acceptance between naive and experienced ants was observed in response to 0.63mM or 0.31mM quinine (17% and 10%. GLMM: z = 1.78, p = 0.3734 and z = 0.10, p = 1, respectively). In contrast to the 10-second acceptance results, no difference in the 90-second acceptance scores (consumption) was found between the groups for any of the quinine concentrations. Specifically:

- 22.5%: 1M sucrose + 1.25mM quinine (GLMM: z = 2.11, p = 0.1755),
- 30%: 1M sucrose + 0.94mM quinine (GLMM: z = 0.09, p = 1),
- 9%: 1M sucrose + 0.63mM quinine (GLMM: z = 1.23, p = 1),
- 7%: 1M sucrose + 0.31mM quinine (GLMM: z = 0.08, p = 1),
- 2%: pure 1M sucrose (GLMM: z = 0.58, p = 1).

**Figure 2:**
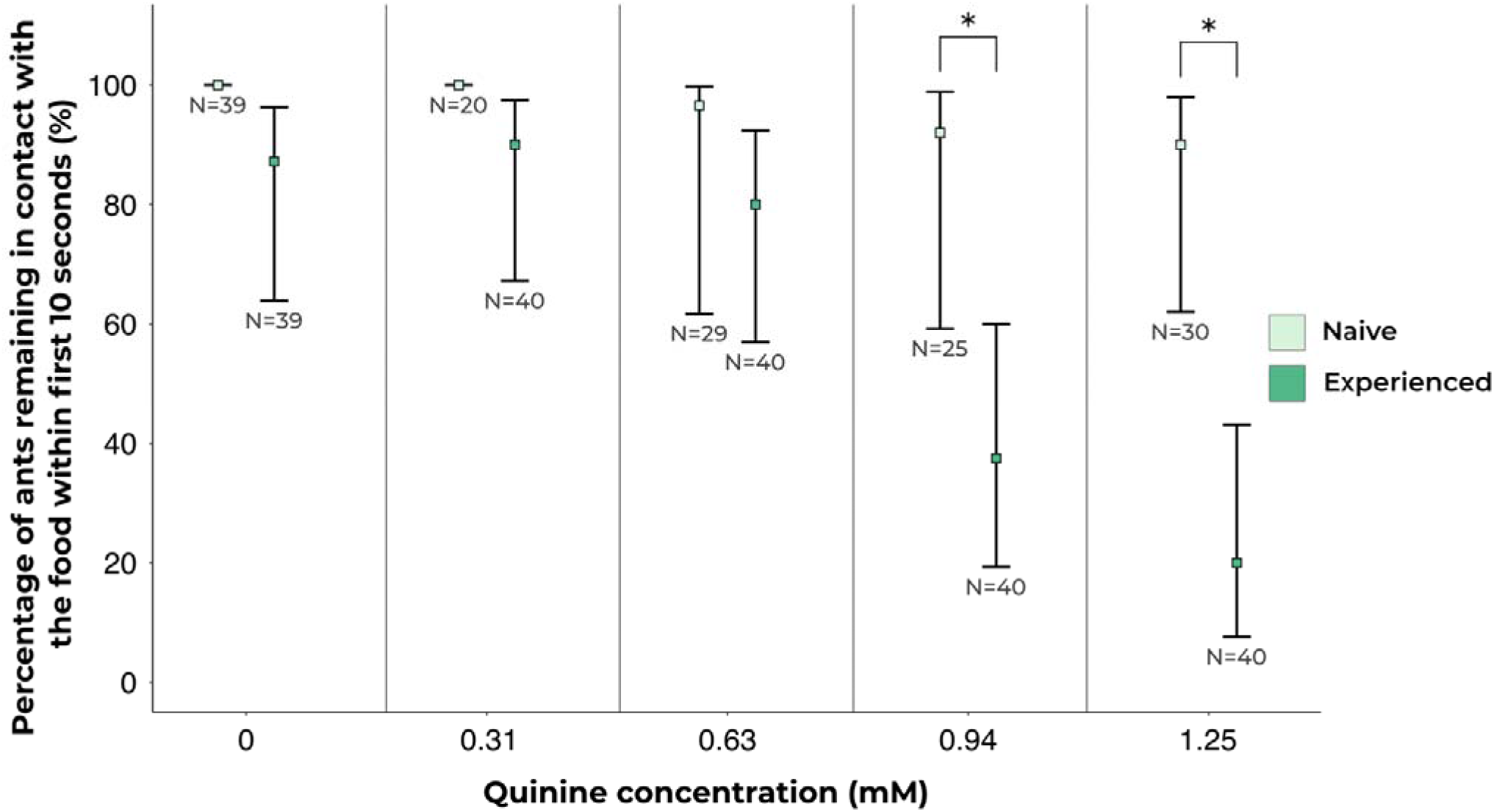
Comparison of acceptance between naive and experienced ants: Naive ants consistently accepted the presented food solutions without showing aversion, while experienced ants, tested on a comparative setup, exhibited aversion to sucrose solutions containing 0.94mM or 1.25mM quinine. Squares represent estimated marginal means obtained from beta regression models, whilst error bars represent the 95% confidence intervals associated with each estimate. Asterisks (*) denote pairs with statistical differences in preference.

On average, five ants were tested by one experimenter per hour.

### Experiment 2 – Quantifying ant acceptance using the dual-feeder method

#### Avoidance of quinine-containing solutions

In this experiment, we evaluated the efficacy of our dual-feeder method by investigating the responses of ants to sucrose solutions infused with varying levels of quinine to assess bitterness rejection. The key measure explored was the proportion of time spent drinking from the first choice feeder solution, because ants which fully accept a liquid food source tend to remain feeding at it until satiated.

A slight right-side bias was observed across trials (62.7% of ants encountered the right well first, betareg: Chisq: 11.59, p = 0.0007). However, there was no effect of quinine side on the proportion of time spent by ants feeding from the test substance, suggesting that the observed right-side bias was not linked to sensory lateralization and did not confound the results (Betareg: Chisq: 0.479, p = 0.488).

Ants significantly preferred solutions with no quinine, spending a greater proportion of time on the first choice feeder when it offered 1M sucrose solution compared to 1M sucrose solution with 0.63mM quinine (20.3%. Betareg: Chisq = 12.79, p = 0.0003). Additionally, ants showed a significant preference for 1M sucrose solution compared to 1M sucrose solution with 0.31mM quinine (16.8%. Betareg: Chisq = 15.25, p = 0.0001). However, the comparison between 1M sucrose solution and 1M sucrose solution with 0.156mM quinine did not show a significant difference (5%. Betareg: Chisq = 0.89, p = 0.344). For a visual representation of the ants’ preference for bitterness levels, see Figure 3. On average, 12 ants were tested by one experimenter per hour.

**Figure 3:**
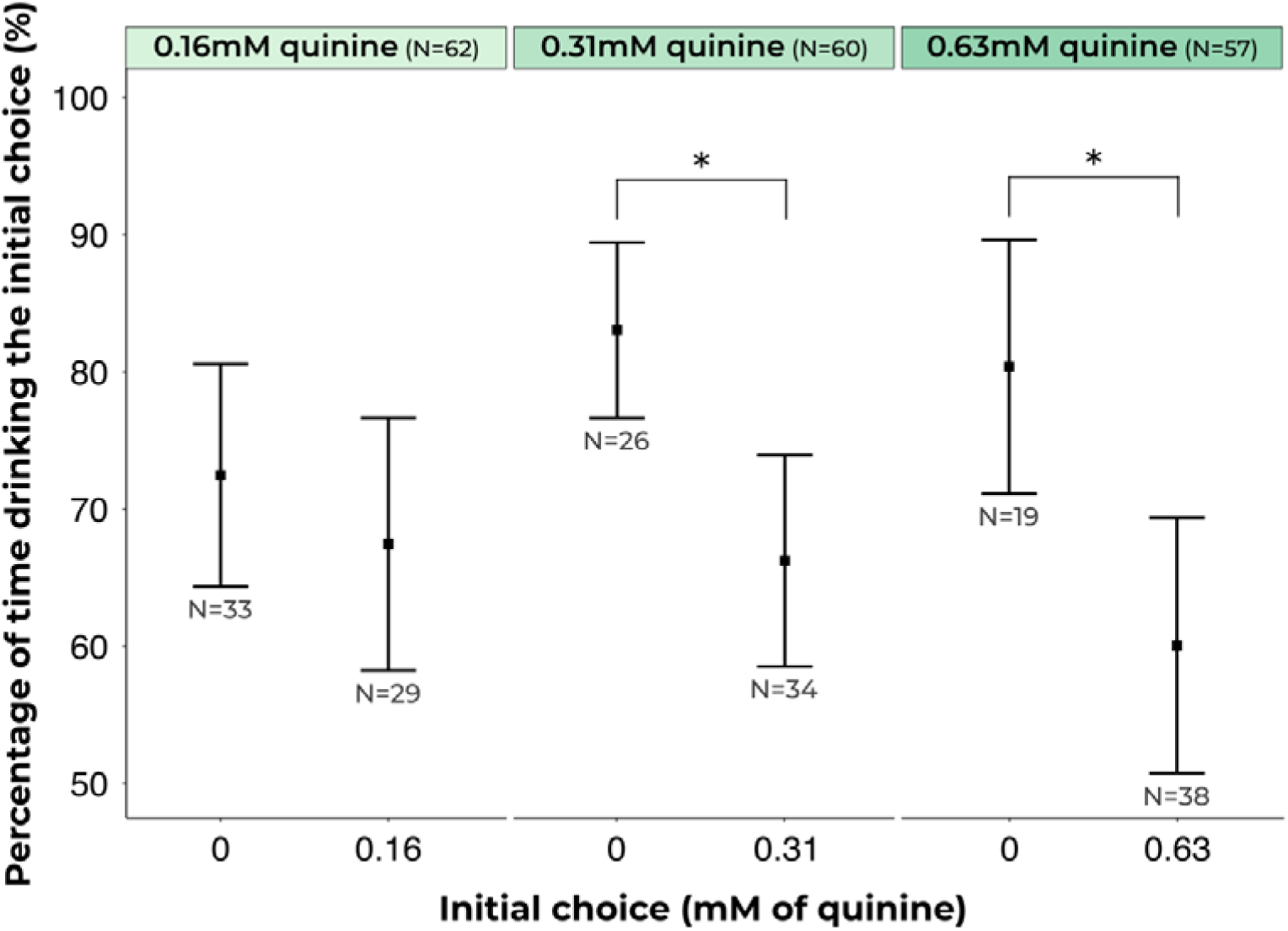
Drinking behaviour across different quinine concentrations and pure sucrose: Ants exhibited a strong preference for pure sucrose solutions (0mM quinine) over sucrose solutions containing quinine (0.63mM and 0.31mM), except for the lowest (0.16mM). Squares represent estimated marginal means obtained from beta regression models, whilst error bars represent the 95% confidence intervals associated with each estimate. Asterisks (*) denote pairs with statistical differences in drinking behaviour.

#### Preference for higher molarity sucrose solutions

Ants significantly preferred higher concentration sucrose solutions, spending a greater proportion of time on the first choice feeder when it was 1M sucrose solution compared to 0.5M sucrose solution (15.8%. Betareg: Chisq = 18.37, p < 0.0001) and also when compared to 0.75M sucrose solution (19.4%. Betareg: Chisq = 12.60, p = 0.0004, see Figure 4). The comparison of 1M sucrose solution to another 1M sucrose solution, serving as a negative control, did not yield differences in preference (1.2%. Betareg: Chisq = 0.24, p = 0.6274). On average, 12 ants were tested by one experimenter per hour.

**Figure 4:**
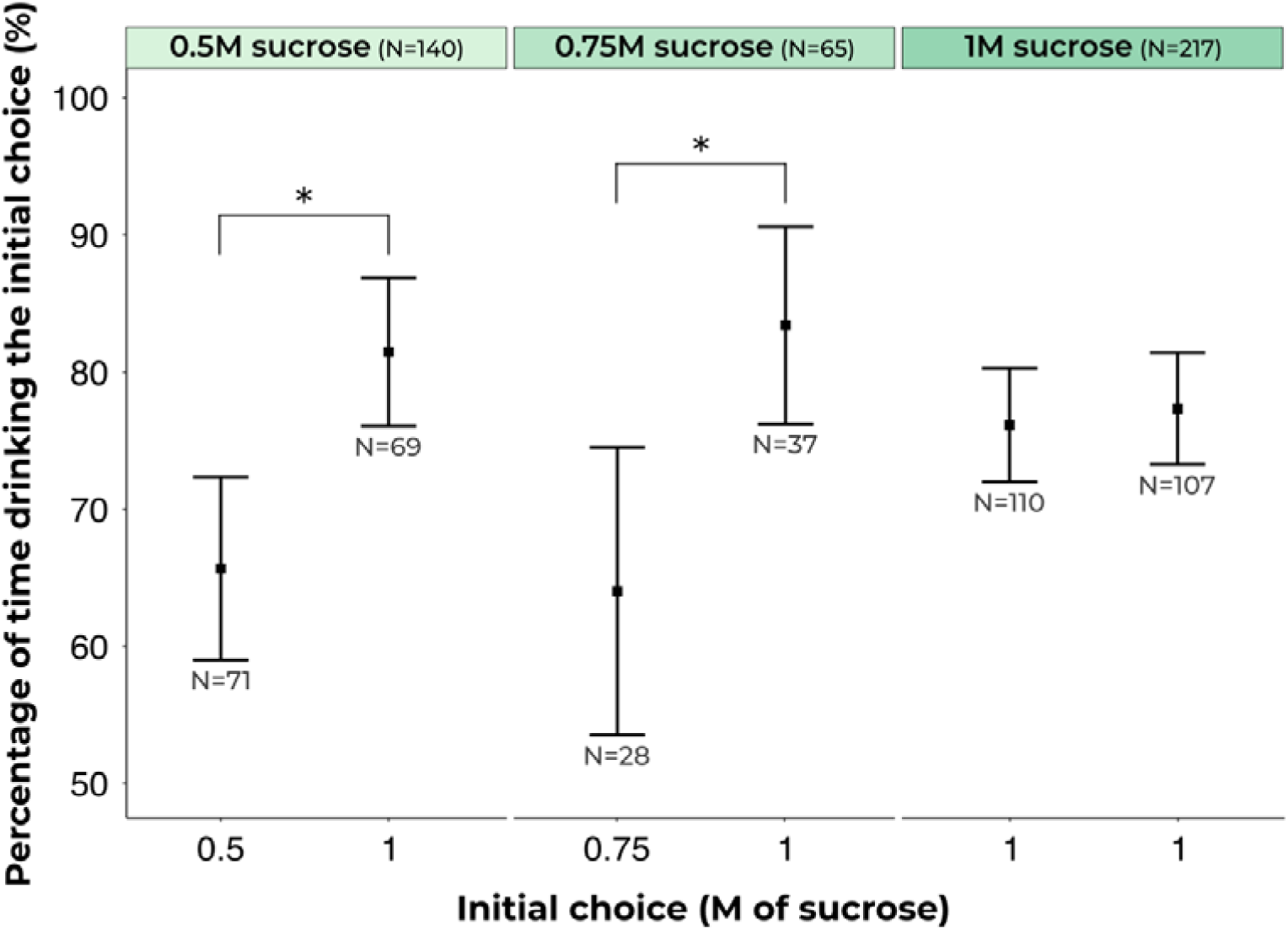
Drinking behaviour across different sucrose concentrations: The figure shows that higher molarity solutions were preferred, while two identical 1M sucrose solutions did not show a significant difference in consumption. The squares represent the estimated marginal means obtained from beta regression analysis, while the error bars indicate the confidence intervals derived from the same regression. Asterisks (*) denote pairs with statistical differences in drinking behaviour.

## Discussion

We demonstrated that comparative evaluation is a more sensitive approach to quantifying ant feeding preference than absolute food acceptance. Our newly developed simultaneously comparative dual-feeder method is faster and easier to use than the alternative sequential comparative preference method. It also avoids triggering neophobia or sequential exposure effects, which have all been reported in ants.^36,46^ Perhaps most importantly, it seems to be more sensitive as well, able to detect preference at a lower level of quinine than the sequential method. Using the dual-feeder method, we demonstrated that ants are capable of differentiating between various levels of sweetness in sucrose solutions and bitterness in sucrose-quinine solutions. The dual-feeder method seems to be especially sensitive to differentiating food sources containing small amounts of disliked compounds.

The marked contrast between the response of naive ants to the moderately bitter solutions with those of ants expecting non-bitter solutions demonstrates that comparative preference evaluation is much more sensitive than absolute acceptance evaluation. However, at moderately low levels of bitterness (0.31mM quinine) no difference between naïve and experienced ants was observed, although data from the dual-feeder method demonstrates that they can, indeed, perceive this level of quinine. One possible reason why we see this difference between the preference testing approaches is that ants weigh the opportunity costs of investing time and energy searching for the previous food source against the known benefits of the present one. Furthermore, ants in the sequential comparative approach might receive feedback from their colony through trophallaxis as indicated in Wagner et al.^46^ Another possibility is that direct evaluation allows the ants to detect smaller differences than evaluations which require a quality stored in the ant’s memory to be compared to those currently being directly sensed.

Interestingly, the same, relatively high, proportion of ants drank to satiation (visible distention of the abdomen after 90 seconds of drinking) in all bitterness levels in the non-comparative and sequential tests. This may be a result of the constrained nature of the experimental design: having potentially rejected the second food source within the first 10 seconds, ants may have spent some time attempting to relocate the initial food source, only to discover that it was no longer available. They may have then decided to accept the only available food source. It is quite possible that, had the ant encountered the less preferred food in an unrestricted area, it would have wandered off without consuming the food.

While clearly more sensitive than the simple acceptance test, the sequential preference testing approach suffers from a number of potential confounds. Firstly, ants have been shown to undervalue unexpected food flavours, even if those flavours are not in themselves distasteful.^36^ Ants also prefer the option they are trained on first.^45^ It is thus impossible to be certain if a reduction in acceptance towards the test food is due to a true dislike of the test food, an expectation disconfirmation effect, or simply because the first experienced food is always preferred. Furthermore, the tested ant must first return to the nest to unload from the first visit. This not only makes the procedure take longer, it also introduces food to the other nestmates, which may change their behaviour if they are subsequently tested.^46–51^ To ensure the testing of truly naive ants, the colonies must be split into donor and recipient fragments, with tested ants only coming from donor fragments and only returning to recipient fragments. This doubles the number of colony boxes needed. Moreover, ants returning to a nest which is not their home nest tend to take longer to unload their crop, and are less likely to return to foraging (TW, TJC, Pers. Obs.). A further issue with the sequential testing approach is that ants may detect a difference between the training (first) and test (second) food, but choose not to reject it, because they consider the cost of rejecting it and going onwards to search for the better test food too high. Finally, this approach requires marking of ants, which may stress the ants, and may be difficult for ‘nervous’ species, such as the invasive Longhorn and Yellow Crazy ants (*Paratrechina longicornis* (Latreille, 1802), *Anoplolepis gracilipes* (Smith, 1857). Given the lack of difference in response between the 1^st^ and 2^nd^ presented food in the control test, marking seemed to not influence acceptance in the current study.

The comparative sequential preference approach and our simultaneous comparative dual-feeder support each other in their results. Nevertheless, we think that the simultaneous comparative approach is superior. The dual-feeder has proven to have higher sensitivity than the sequential approach. This may be because the simultaneous evaluation reduces the impact of other factors, as mentioned above. Our behavioural tests thus demonstrate that *L. humile* can perceive quinine in sucrose solutions at concentrations at least as low as 0.31mM, and prefer pure sucrose solution. Only at very low quinine concentrations (0.16mM) could we detect no preference for the pure sucrose solution. This may be because the ants could not perceive the quinine – either because it is absolutely below their perception threshold, or because at these low levels perception of the quinine is masked by the sweetness of the solution. A potential masking effect means that the detection threshold we measured may be an underestimate. Masking of flavours is poorly studied in insect, but does occur: for example, in *Manduca sexta* caterpillars, bitterness from caffeine was successfully masked by some sugars, including sucrose.^52^ Alternatively, the ants may have perceived the quinine but not have found it aversive at such low levels.

The dual-feeder approach can also be used to differentiate preferences between attractive solutions with no aversive components. Testing with the dual-feeder method is rapid, enabling the examination of approximately 12 ants per hour for a minimally-trained experimenter, compared to sequential testing at 5 ants per hour. Moreover, no marking or returning to the nest is needed, thereby avoiding many of the confounding issues mentioned above. In cafeteria tests, ants are also presented with multiple choices of different food sources simultaneously. However, contrary to our dual feeder, most individual ants in a cafeteria test may not encounter more than one food source, thus hindering their ability to perform a comparative evaluation. When running cafeteria experiments, often an entire colony or colony fragment is given access to the food sources. Due to recruitment and other positive-feedback processes, individual ant choice is not independent, and the replication unit of cafeteria experiments is thus the colony, necessitating a much higher number of colonies, and ants, to achieve robust results.

Our dual-feeder method was tested with *Linepithema humile*, but we believe it has potential for application across a variety of ant species. Minor adjustments to the feeder size would likely be sufficient to accommodate a wide range of species - especially invasive and pest ants which tend to tend homopterans and rely heavily on carbohydrates. More broadly, the concept of simultaneous comparative preference testing could and should be adapted to alternative feeder designs tailored to the specific morphology or feeding habits of other target organisms.

The dual-feeder system boasts high sensitivity and a high throughput. However, it is important to acknowledge that it has certain limitations. The potential for lateralization in antennal function could influence the ants’ perception of the solutions presented. Lateralization has been demonstrated in *Apis mellifera* (Linnaeus, 1758), where bees respond more sensitively to odours when trained through their right antenna.^53–55^ Similar findings have been found in the bumblebee *Bombus terrestris* (Linnaeus, 1758).^56^ Indeed, a right antenna lateralization has also been observed in ants in the context of trophallaxis.^57^ It may be that placing the test solution in the right well of the dual feeder may result in higher sensitivity. However, this was mitigated by alternating the placement of different solutions (sucrose and sucrose-quinine mixtures) on the left and right sides of the dual-feeder, minimizes the likelihood that any innate lateralization in antennal use would significantly impact the outcomes. Moreover, we found that that quinine was not rejected more often when placed on the right side of the feeder compared to the left, speaking against a sensitivity lateralisation effect in this experiment. If lateralisation were to be found in other species, it could indeed be harnessed by placing the test food on the more sensitive side.

Another limitation of the dual feeder method is that while positive results are indicative of detection as well as preference, negative results indicate either no preference or no detection. Finally, in natural foraging scenarios, ants are likely to encounter conditions more akin to a cafeteria test or sequential comparisons, where choices are influenced by prior experiences and interactions with nestmates, rather than the simultaneous presentation of options as in our dual-feeder setup. While our method demonstrates heightened sensitivity, it’s important to note that the detection and disfavouring of a substance under these controlled conditions may not mirror absolute rejection in the wild. On the other hand, in natural settings, ants might exhibit reduced recruitment to less preferred sources, an aspect that our study did not directly investigate. We note that acceptance scores and recruitment intensity tend to correlate.^58^

The dual-feeder method we propose is conceived as an early-stage laboratory assay, designed to sensitively detect aversion to, and preference for, bait formulations and individual additives. However, translating laboratory findings to the field is rarely straightforward. In the field, environmental factors such as temperature fluctuations, humidity, and the availability of diverse food sources could influence ant behaviour and preferences in ways that are not fully captured in a lab setting. For example, warmer food may enhance both appetitive and aversive stimuli in a bait.^59^ In addition, social interactions within the colony, such as trophallaxis and recruitment dynamics could alter individual preferences and affect the overall colony’s response to bait formulations.^48, 59^ In natural settings, ants may be more selective or exhibit different levels of recruitment depending on competition from other food sources or the presence of nestmates.^22,60^ While the approach we propose could be used to systematically examine the influence of abiotic and biotic factors on food preference, they cannot replace field trials. Rather, we hope our approach can serve as a valuable tool in early-stage testing, increasing the success rate of formulations which are taken on to field trials. In a separate study, we demonstrated the efficacy of the dual-feeder method for answering control-related questions, by examining the effect of three major toxicants, and egg protein, on food acceptance.^62^

It is important to note that different ant species have different sensitivities to aversive substances, and that *Linepithema humile* is especially tolerant of bitter substances (in prep). Other ant species may show much higher sensitivities than those described here. Creating effective and species-specific baits is challenging due to the diversity of ant preferences and the complexity of integrating repellent substances without reducing bait attractiveness.^22,58^ Nyamukwondiwa and Addision^64^ note that developing attractive baits while including deterrents is a major bottleneck in the development of effective ant control strategies. Our dual-feeder method offers itself as a powerful approach for testing potential bait solutions quickly and effectively. With the continued emergence and spread of invasive ants, innovative and sustainable approaches in bait formulation have become a top priority.^60,61^ We anticipate that the dual-feeder method we report here can contribute to these ongoing efforts. Moreover, by simply scaling the size of the apparatus, this approach could easily be deployed for testing a range ant species.

## ACKNOWLEDGEMENTS

We would like to thank Silvia Abril for providing ant colonies. Thomas Wagner and Henrique Galante were supported by an ERC starter grant (Cognitive control: 948181) to Tomer J. Czaczkes. Tomer J. Czaczkes was supported by a Heisenberg fellowship from the Deutsche Forschungsgemeinschaft (CZ 237/4-1). Many thanks to Cosmina Werneke, Laura Wögler, Leonie Dechant and Nick Wolter for conducting part of the experiments.

## DATA AVAILABILITY STATEMENT

The data that supports the findings of this study are available online from https://doi.org/10.5281/zenodo.10953784.

## AUTHOR CONTRIBUTIONS

1. T. Wagner: Conceptualization, Methodology, Investigation, Writing - Original Draft, Writing - Review & Editing, Supervision. H. Galante: Software, Validation, Formal Analysis, Data Curation, Writing - Review & Editing, Visualization. T. J. Czaczkes: Conceptualization, Methodology, Resources, Writing - Review & Editing, Supervision, Project administration, Funding acquisition.

